# Basal Forebrain Deep Brain Stimulation Impacts the Regulation of Extracellular Vesicle Related Proteins in the Rat Brain

**DOI:** 10.1101/374256

**Authors:** Wenxue Li, Laura Lozano Montes, Jayakrishnan Nair, Marta Dimanico, Reza Mazloum, Zehan Hu, Brigitte Scolari, Jörn Dengjel, Franziska Theilig, Michael Harvey, Xiaozhe Zhang, Gregor Rainer

**Author notes:** Raw data are available via ProteomeXchange with identifier PXD008426, Username Password: f0df0Hsz.

## Abstract

Extracellular vesicle (EV) signaling has attracted considerable attention in recent years because EVs play a key role in long distance cellular communication functions. EV studies have begun to reveal aspects of physiological and physiopathological regulation in numerous applications, although many areas remain to date largely unexplored. Deep brain stimulation (DBS) has shown remarkable therapeutic benefits of patients with neuropsychiatric disorders, but despite of the long and successful history of use, the mechanisms of action on neural ensemble activity are not yet fully understood. Here we explore how DBS of the basal forebrain impacts EV signaling in the rat brain. We employed differential centrifugations to isolate the EVs prefrontal cortex (PFC), hippocampus and striatum. We then performed quantitative analysis of EV-associated proteins using an MS-based proteomics method. We identified a considerable number of EV-associated proteins are modulated by DBS in three brain regions, some of which have been previously linked with central nervous system disorders. Particularly, neurofilament proteins NFL and NFM were both significantly changed in EVs of PFC, hippocampus and striatum after DBS stimulation compared with controls. The SOD1 protein, associated previously with neurodegenerative diseases, was significantly increased only in PFC. Our study is the first, to our knowledge, to use EV protein analysis to examine DBS effects on brain physiological regulation. Our findings open an entirely new perspective on brain area specific DBS effects.

## Introduction

Extracellular vesicles (EVs), which are most prominently involved in shuttling reciprocal signals between myelinating glia and neurons, contribute to many aspects of central nervous system (CNS) development and function, including regulation of synaptic communication, synaptic strength, and synapse assembly and plasticity (1–3). EVs can selectively shuttle cytosolic and membrane proteins, as well as RNA and lipids, across cells to achieve an exchange of signaling molecules (4, 5). For example, cultured neurons from embryonic and mature mammalian neural tissue release exosome-like EVs after stimulation with depolarization or alternative means to induce excitation, such as application of Ca^2+^ ionophores, GABA receptor blockers or Glutamate receptor agonists α-amino-3-hydroxy-5-methyl-4-isoxazole-propionic acid (AMPA) or *N*-methyl-D-aspartate (NMDA) (6, 7). EVs have also been identified as important factors in the progression of neurodegenerative diseases (2), such that EVs have been implicated in the regulation of amyloid-ß protein (8–11), tau protein (12–14), alpha-synuclein (15–18), TAR DNA-binding protein 43 (TDP-43) (19–21) and leucine-rich repeat serine/threonine kinase 2 (LRRK2) (22, 23). Some of these proteins are considered to be potential biomarkers for neurodegenerative diseases such as Alzheimer disease (AD), Parkinson disease and amyotrophic lateral sclerosis (ALS). For example, alpha-synuclein, which is packaged into EVs for extracellular transport and released into extracellular space, has been closely linked to the pathogenesis of PD (16). Additional evidence suggests that EVs play a role in the sequestration of toxic oligomers and in the release of oligomers from protein aggregates (24–26). The loss of functional aggregated proteins can interference with axonal transport and lead to proteasomal inhibition, synaptic toxicity and endoplasmic reticulum stress (27–32). While the mechanisms that contribute to neurodegenerative diseases are diverse, protein aggregates are widely considered to have a central role (2), and EVs are thus likely candidates as vehicles of propagation of neurodegenerative disease and the cell- or region-specific cargos that EVs carry give them the potential to harbor disease-specific molecular signature.

The modulation of brain activity by way of direct electrical stimulation has a long history, with early studies more than a century ago showing that motor cortex stimulation in dogs can elicit limb movements (33). In recent years, deep brain stimulation (DBS) in specific brain regions has provided therapeutic benefits for otherwise treatment-resistant disorders such as chronic pain (34, 35), Parkinson’s disease (36, 37), essential tremor (38, 39), dystonia (40, 41), major depressive disorder (MDD) (42), obsessive-compulsive disorder (OCD) (43), Tourette’s syndrome (44–46), addiction (47) and obesity (48). Furthermore, for a number of these disorders DBS has been approved for clinical therapy. Advantages of DBS include the direct and spatially specific modulation of brain activity as well as the controlled manner of stimulation and reversibility of effects. Despite the long history of DBS, its underlying principles and mechanisms remain incompletely understood (49–51). A large number of studies have reported the release of neurotransmitters induced by DBS in brain areas that are relevant for movement disorders. Initial studies on rats showed that the subthalamic nucleus (STN) stimulation increase extracellular glutamate levels in the STN, Globus pallidus (GPi) and substantia nigra pars reticulata (SNpr) (52). In anaesthetized 6-OHDA-lesioned rats, STN stimulation caused GABA increasing in the SNpr (53), while another study showed the STN stimulation may differentially regulate both glutamate and GABA release (54). Parkinson’s disease patients showed increase of cyclic GMP in the GPi instead of increasing of extracellular glutamate (55). Based on this evidence, we hypothesized that DBS might regulate EVs containing signaling molecules in a brain region specific manner.

The basal forebrain (BF) is considered to be a major cholinergic hub of the central nervous system. It consists of several structures that send cholinergic projections to the cortex, hippocampus and other brain regions where they play an important role in various cognitive functions including learning, memory, formation and attention. The Nucleus basalis magnocellularis (NBM) is part of this structure, projecting mainly to the neocortex, amygdala, and thalamus (56, 57). Previous studies showed that the NBM has important role in the neural activity in learning and memory (58–62), such that for example NBM lesions affect the acquisition of spatial memory and produce a severe selective disturbance in recent memories (63, 64) and electrical NBM stimulation improves cognitive functions of patients suffering from dementia (65). Effects of NBM electrical stimulation have been demonstrated also at the level of neural activation patterns in cortex, such that for example BF activation robustly enhances contrast sensitivity in visual cortex (VC), an effect that likely involves cholinergic modulation but also BF GABAergic projections to VC (66). The effects of DBS at the level of physiological signaling cascades has been much less investigated.

Here we hypothesize that BF simulation may also exert a profound influence on signaling pathways within the BF and at BF projection brain regions. We employed differential centrifugations for isolation of EVs from PFC, hippocampus and striatum brain regions after unilateral electrical stimulation in the NBM of the rat brain, with the other non-stimulated brain hemisphere serving as control. Such an internal control allows us to reduce data variability across animals and improves the accuracy of results. In addition, we employed the dimethyl label method for relative quantification on levels of proteins and peptides. We observed a large number of significantly modulated proteins in brain regions of the stimulated hemisphere compared with the control hemisphere. Using functional analysis with the DAVID algorithm, we linked these identified EV-proteins to oxidative stress, DNA damage, nervous system development and signaling processes.

## EXPERIMENTAL SECTION

### Materials

LC-MS grade acetonitrile and formic acid were purchased from Fisher and Fluka, respectively. Acetic acid was purchased from Fluka (Buchs, Switzerland). Pure water was prepared by GenPure system (TKA, Niederelbert, Germany). Siliconized micro centrifuge tubes were purchased from Eppendorf (Hamburg, Germany). Formaldehyde (CH_2_O, 37%, v/v, cat. no. 252549), sodium cyanoborohydride (NaBH_3_CN, cat. no. 156159) and triethylammonium bicarbonate buffer (TEAB, cat. No. T7408) were purchased from Sigma. Sodium cyanoborohydride (NaBD3CN, 96% D, cat. no. 190020) and formaldehyde (^13^CD_2_0, 20%, 99% ^13^C, 98% D, cat. no. 596388) was purchased from Cambridge Isotope Laboratory, Inc (Tewksbury, UK).

### DBS animals

DBS is known to have an excitatory or inhibitory effect depending on stimulation frequency stimulation, although results tend to vary as a function of stimulation parameters and brain region under study (67–69), and biphasic cathode-first stimulation tends to be more effective in eliciting functional responses (70–72). In the present study, we tested different stimulation parameter combinations and selected biphasic pulse trains with 20 Hz frequency, ±8 V amplitude, 100μs pulse duration, 500 ms pulse train duration and 10s pulse trains intervals. This stimulation protocol was applied during two consecutive days with three stimulation sessions per day lasting 1h each with a 1h pause between sessions. BF electrodes were implanted bilaterally, but only one hemisphere was stimulated in each animal with the other hemisphere serving as a control. A total of n=6 animals were used in this study.

### EVs Isolation and Sample Preparation

The EVs isolation protocol was modified according to a previous report (73). In order to obtain enough analyte for the procedures, samples from two animals were pooled together as one biological replicate. Brain tissue was dissected and treated with 20 units/ml papain with Hibernate A (Gibco) for 15 min at 37 °C. It was then gently washed with another two volumes cold Hibernate A and then sequentially centrifuged 3000 g for 20 min and then filtered through a 40 μm mesh filter (BD Bioscience) and a 1 μm syringe filter (Acrodisc, Sigma). EVs isolation centrifuge steps were modified as described previously (74). Briefly, the filtrate was sequentially centrifuged at 5000g for 20 min at 4 °C to discard small debris. The supernatant was centrifuged at 20,000g for 1 h at 4 °C. The pellets were washed with cold PBS twice collecting pellets as microvesicles, and supernatants were further performed ultracentrifuge 200,000 g for 2 h at 4 °C. The pellets were washed with PBS twice collected as exosomes. The isolated EVs were incubated with 100 μL 8 M urea (pH=7.4) containing Cocktail proteinase inhibitors for 30 min in the ice after vortex properly. 50 mM DTT and IAA were first used for disulfide bond reduction and alkylation as described, respectively. Urea concentration was diluted to below 1 M for trypsin (Roche) digestion at 37 °C for overnight. The dimethyl labeling was performed after desalting with C18 StageTip (75). The dried samples were resolved with 100 mM TEAB first and subsequently, add 5 μL of 4% (v/v) CH_2_O and ^13^CD_2_O, 5 μL of 0.6 M NaBH_3_CN and NaBD_3_CN to the samples for “light” and “heavy” dimethyl labeling, respectively. The labeling reaction was put in a working fume hood for 1 hat room temperature. Quench the labeling reaction by adding 10 μL of 1% (v/v) ammonia solution and then acidify the sample with 20 μL 0.2 % formic acid. The differential labeled samples were mixed first and desalted with C18 StageTips (75) again, which pre-washed with 200 μL 0.2 % acetonitrile and 200 μL 0.2% formic acid in water twice, respectively. Finally, 200 μL of 80% acetonitrile containing 0.2% formic acid was used for eluting peptides and then directly analysis by nanoLC-MS with three replicates. For each peptide sample, three technique replicates were performed.

### Transmission Electromicroscopy (TEM)

Microvesicle and exosome fractions were enriched with above centrifugation steps and then fixed in 2% paraformaldehyde/ 2.5% glutaraldehyde in 0.1M cacodylate buffer solution for 30 minutes, washed and embedded in 2% AGAR/water. Specimens were postfixed in 1% osmium tetroxide in 0.1 M cacodylate buffer for 30 minutes, washed, dehydrated and epoxy resin embedded. 70 nm ultrathin sections were contrasted with uranyl acetate and lead citrate and examined by TEM (Philips CM100).

### LC-MS Data Acquisition

Samples were measured on Orbitrap Discovery (Thermo Fisher Scientific) coupled with UltiMate™ 3000 RSLCnano System (Thermo Fisher Scientific). Analytical column tips with 75 μm inner diameter (New Objective) were self-packed with 1.9 μm Reprosil-Pur C18 AQ (100Å, Dr. Maisch GmbH) to a length of 20 cm. The precolumn (μ-precolumn, C18 Pepmap, 300 μm ID, Thermo Scientific) was used for sample loading with 3 μL/min 100% A (0.2% formic acid in water) in loading pump and then switch to analytical column after 5 min. A gradient of A and B (0.2% formic acid in 80% ACN in water) with increasing organic proportion was used for peptide separation (separation ramp: from 2-30% B within 170 min). The separation flow rate was 300 nL/min and totally 190 min running for each sample. The mass spectrometry was operated in the data-dependent mode and switched automatically between MS (Orbitrap analyzer, resolution 30,000, AGC 1e^6^) and MS/MS (Ion trap analyzer, AGC 1e^4^) with maximum top ten. The fragmentation was performed in CID mode with normalized collision energy 35%. Parent ions with a charge state of z =1 and unassigned charge state were excluded for fragmentation. The mass range of MS was from 400 to 1800 m/z. The other mass spectrometry parameters were as follows: spay voltage 2.5 kV; no sheath and auxiliary gas flow; and ion-transfer tube temperature 200 °C.

### Data analysis and Statistics

All the raw MS data files were processed on MaxQuant software (version 1.6.1.0) (76) for proteins and peptides identification and quantification, and trypsin was specified as digestion for Uniprot rat database (January 2017) searching with a false discovery rate (FDR)<0.01 at level of proteins, peptides, and modifications. DimethLys0 and DimethNter0 were specified as “Light Labels”; DimethLys8 and DimethNter8 were specific as “Heavy Labels”, and 3 maximum labeling for each peptide. Acetylation (protein N-term), phosphorylation (S/T/Y), and oxidation on methionine were selected for the variable PTMs with 3 maximum modifications for each peptide, and carbamidomethyl as fixed modification. The minimum peptide length is seven amino acids, and “match between runs” enabled with a matching time window of 1 min. Proteins and peptides were identified using the Andromeda search engine integrated into the MaxQuant environment (77). Bioinformatics analysis was performed on Perseus(78) and significant A with p-value (FDR 0.05) was used for proteins significantly analysis. A site localization probability of 0.75 was used as cutoff for localization of phosphopeptides sites. The phosphorylation motifs were determined by pLogo program (79). The proteins network was analyzed with STRING(80) and visualized by Cytoscape (81). We used Perseus significance A (78) to determine differential expression of EV proteins in the PFC, hippocampus and striatum of stimulated samples compared with controls.

## Results

EVs, including microvesicles and exosomes, were isolated from rat brain tissue PFC, hippocampus and striatum regions with differential high speed centrifugations and ultrahigh speed centrifugations after incubation with papain and series filtration steps (see methods, Figure 1 and Figure S1) (82). TEM was employed to confirm that excellent EV purity was achieved using this procedure (Figure 1C). After lysis of EVs, dimethyl reagents were used for peptide level stable isotope labeling (83), and mixed peptides were desalted with StageTips (75) and analyzed by liquid chromatography-tandem mass spectrometry (LC-MS/MS) on a high-speed, high-resolution mass spectrometer.

**Figure 1.**
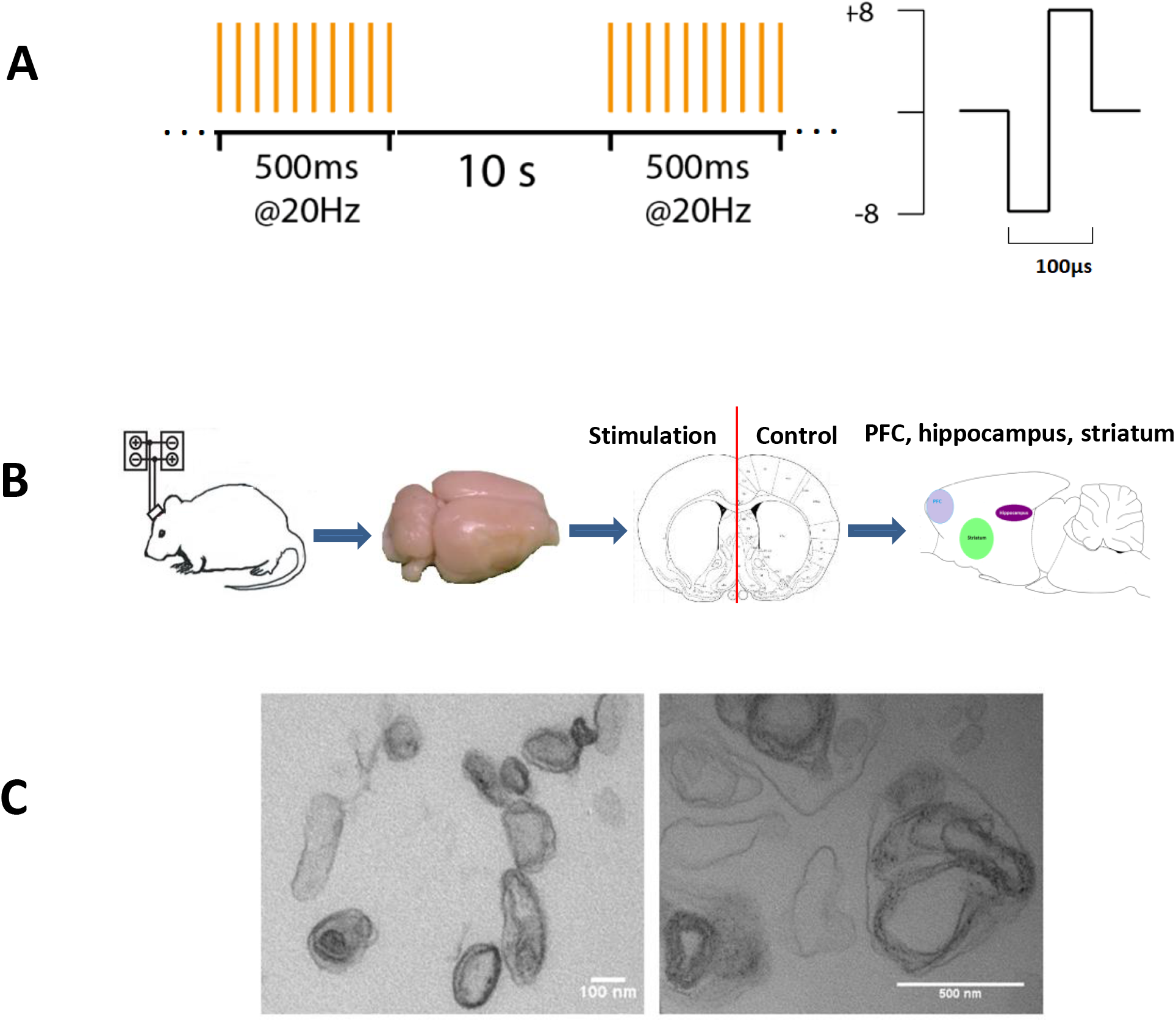
The scheme of stimulation procedure in rat. A, the deep brain stimulation related parameters; B, the workflow of animal experiment; C, the transmission electromicroscopy picture for EVs after isolation, left, exosome, right, microvesicles.

Our strategy allowed us to identify 1037, 634 and 713 EV proteins from PFC, hippocampus and striatum brain regions, which were derived from 1292, 744 and 861 genes, respectively. Gene ontology analysis of the identified EVs proteins indicated overall similar cellular components and biological functions in PFC, hippocampus and striatum (Figure 2A and 2B). In Figure 2C, we illustrate the gene overlap of identified EV proteins from PFC, hippocampus and striatum. Note that the identification ratio tended to be higher for the PFC compared to the other two regions; an effect we attribute to the relatively large tissue sample for PFC. Common EV biomarkers were identified in different brain regions (Figure 2D), such that the EV marker proteins CD9, CD81, CD82 and HSC70 were in fact present in all three brain areas (PFC, hippocampus and striatum), emphasizing the high purity of our EV isolation procedure from tissue samples.

**Figure 2.**
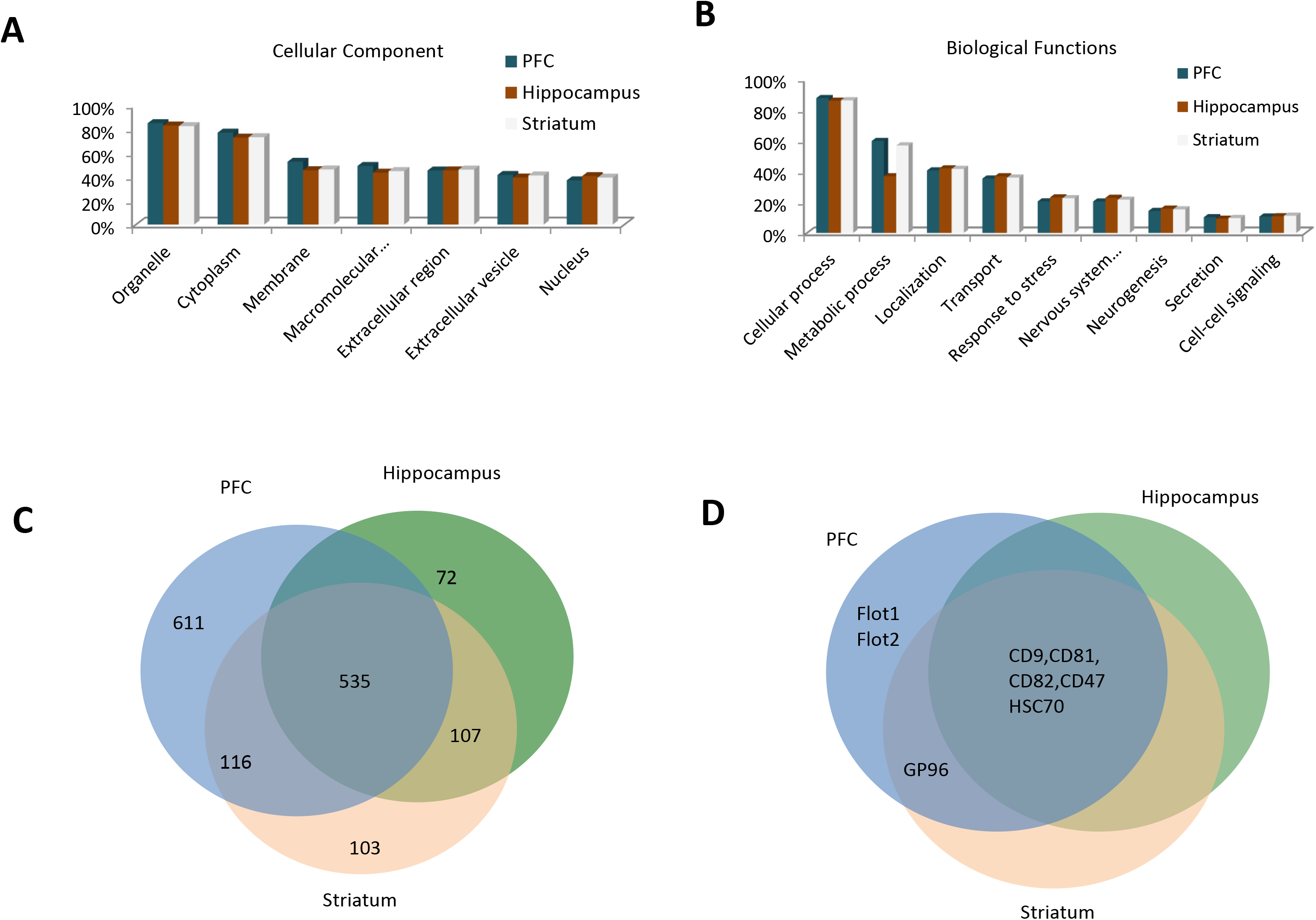
The EV proteins identification from PFC, hippocampus and striatum. A, the cellular component distribution of EV proteins identified from three brain regions; B, the biological function distribution of EV proteins identified from three brain regions; C, the genes overlap of identified EV protein from the three brain regions; D the comparison of EV marker proteins identified from different brain regions.

Based on quantitative statistical analysis, 23, 20 and 24 significantly changed EVs proteins were identified in PFC, hippocampus and striatum respectively. These proteins were derived from 39, 32 and 35 genes respectively. A complete list of all identified and significantly DBS modulated proteins are shown at Supporting Information Table S1. Approximately fifth percentage biological replicate correlation in PFC, hippocampus and striatum has been shown at Figure 3A-C. Since EVs have important communication roles in the CNS, we focused analyses on identified phosphopeptides that result from post-translational modifications of signaling molecules. Note that these analyses were possible despite the fact that no phosphopeptides enrichment was employed in this work. We identified 549, 88 and 552 phosphopeptides respectively in PFC, hippocampus and striatum. The phosphorylation site localization probability was 0.75 and pS/pT/pY site and sequence motif distributions are shown at Figures 3D and Figure S2-5. We observe that the hippocampus exhibits less phosphopeptides identification than other regions, and pY EVs are more common in PFC than hippocampus or striatum. In addition, the hippocampal pT site proportion was significantly increased compared to the other regions. The complete list of identified phosphopeptides were shown at Supporting Information Table S2. All of these disparities may bear relevance to differential brain regions in pathological processes.

**Figure 3.**
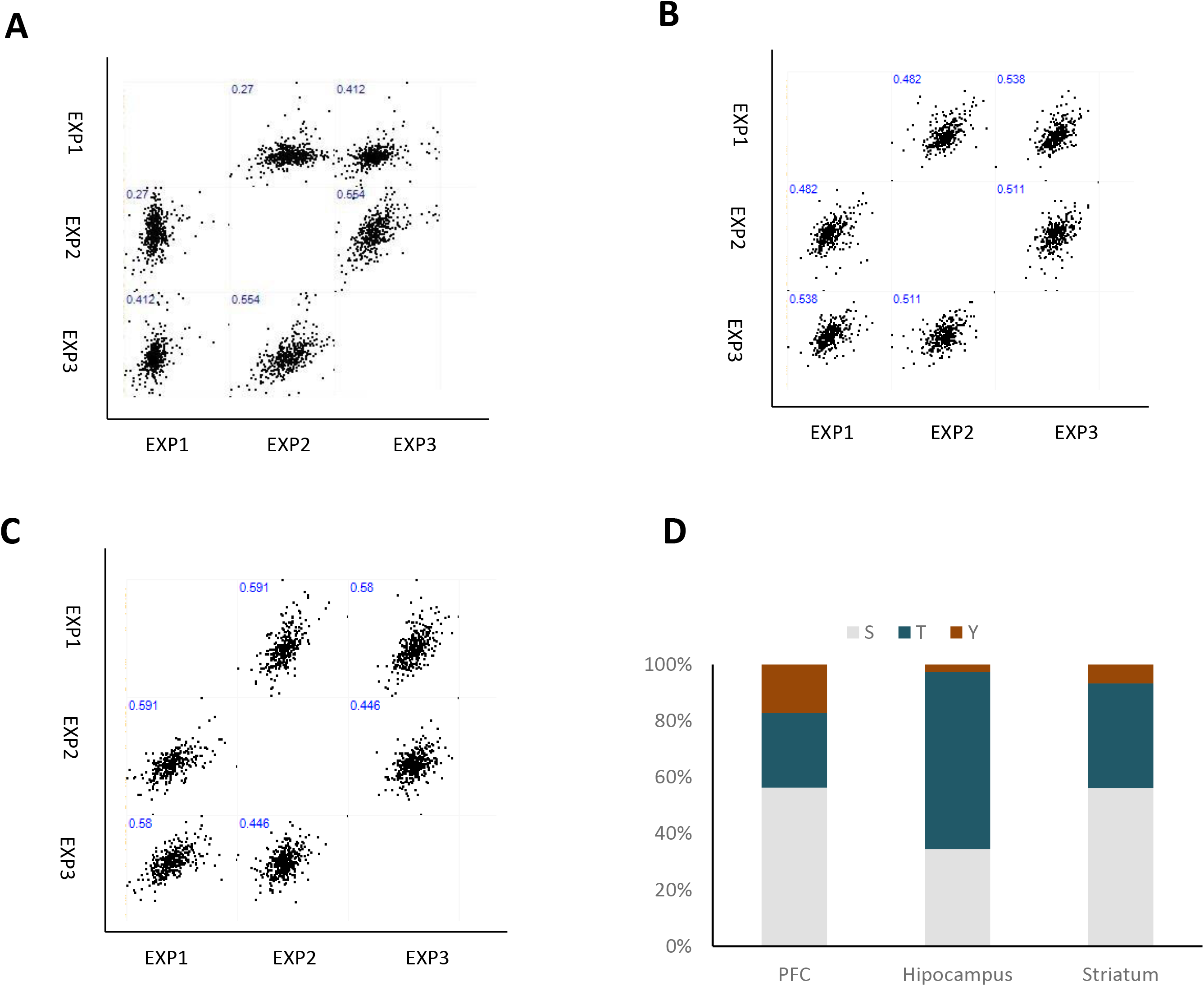
The biological Pearson’s correlation of EV proteins for PFC (A), hippocampus (B) and striatum (C) brain regions. D, distribution of S/T/Y phosphopeptides from EVs in three brain regions.

To better understand the biological roles of differentially expressed EV proteins, we examined the relationship between EV proteins and specific CNS related diseases, using STRING to identify enriched gene ontology categories and signaling networks. In PFC, these significant changed proteins were classified into protein biosynthetic process, oxidative stress, signaling, cytoskeleton and nervous system development processes. For the hippocampus, DNA damage, oxidative stress/nervous system development and signaling related EVs proteins were found to be significantly modulated. The striatum had similar EV protein function classification as hippocampus, with exception of an additional role in protein biosynthetic processes. It is interesting to note that the oxidative stress plays a central role in stimulation caused EV proteins differential expression compared with controls, which is consistent with the previous studies that oxidative stress may be involved in many neurodegenerative diseases(84). Network visualizations of these effects are shown in Figure 4. Focusing on specific proteins, we found that the two NF proteins NFL and NFM were significantly modulated in all three brain regions, but in different directions. NFL and NFM were both downregulated average −5.6 times (−2.0 ~ −14.1) and −10.8 times (−4.2 ~ −23.5) in PFC EVs in stimulated samples compared with control; whereas upregulation both average 2.2 times (1.2 ~ 3.1) and 2.1 times (1.5 ~ 2.7) was observed in hippocampus and striatum (Figure 5A), respectively. Protein NFH was also identified but levels for this protein did not modulate significantly excepting in PFC EVs (reduced average −8.4 times (−3.6 ~ −21.3)). These findings are consistent with previous studies suggesting that NF proteins may be involved in neurodegeneration (85). We proceeded to summarize NF proteins subunits that have different regulation in different brain regions in neurodegenerative diseases and others CNS related disease studies, such as Schizophrenia and MDD (86–88) as illustrated in Figure 5B and 5C. These data support the idea that the EVs proteome is a sensitive indicator for physiological regulation and that quantitative analysis of EVs proteome can be disease specific.

**Figure 4.**
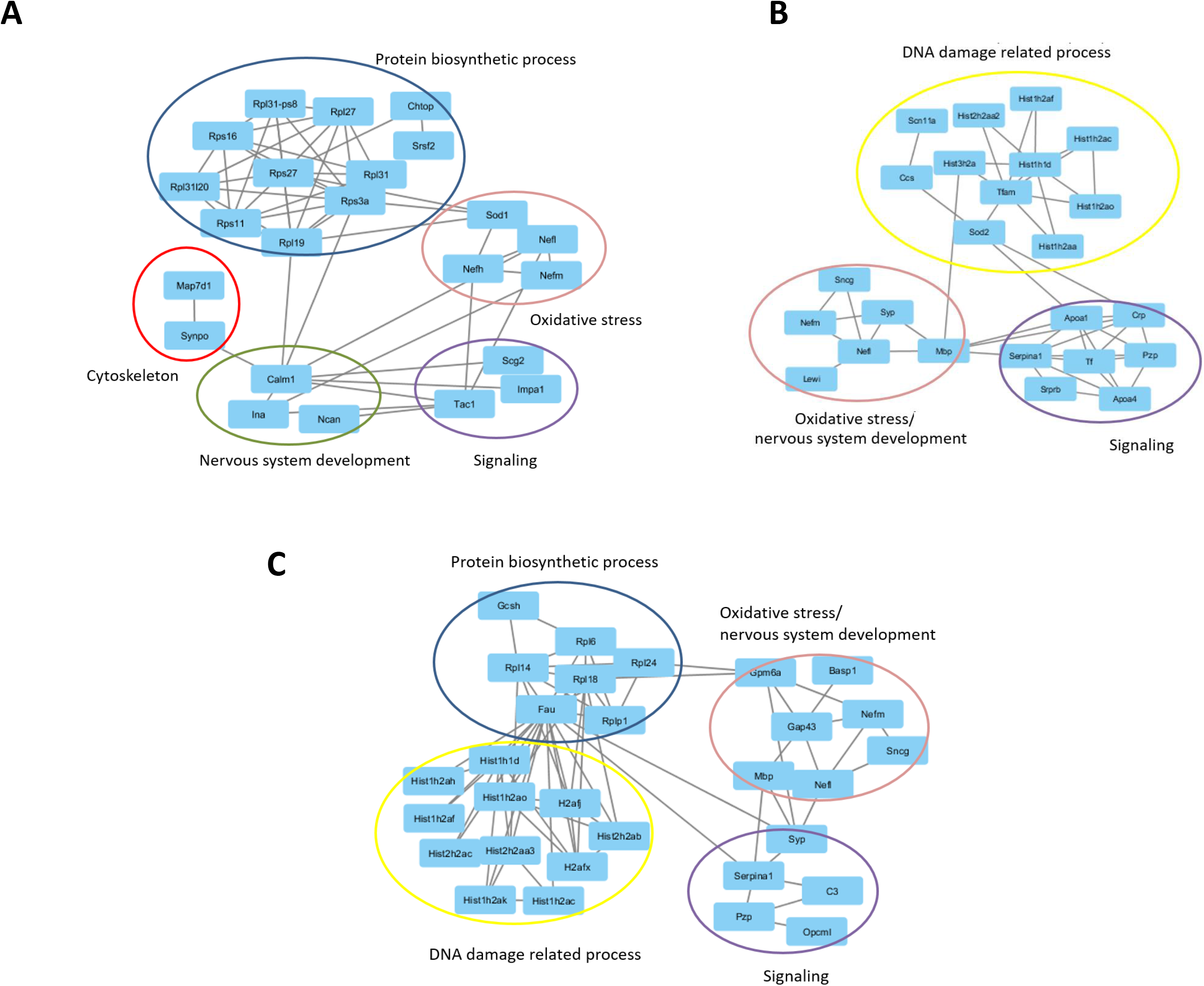
The network analysis for significant changed EV proteins from PFC (A), hippocampus (B) and striatum (C).

**Figure 5.**
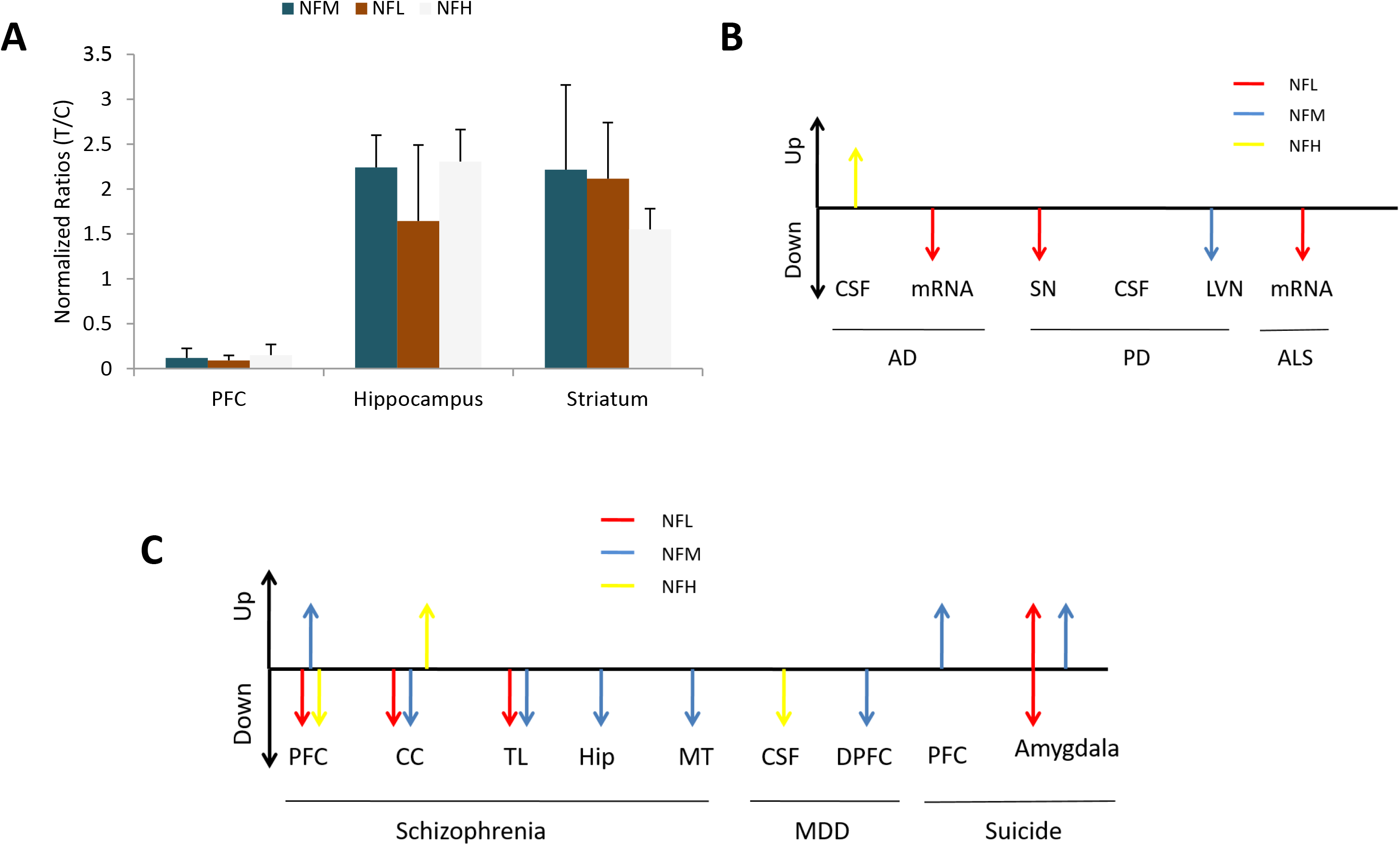
The neurofilament (NF) proteins regulation direction comparison with literatures. A, the different alterations of NF proteins in PFC, hippocampus and striatum with the brain stimulation; B, the NFL, NFM, NFH proteins different regulation in neurodegenerative diseases according to literatures. C, the NFL, NFM, NFH proteins different regulation in Schizophrenia, MDD and suicide. NFL, neurofilament light polypeptide; NFM, neurofilament medium polypeptide; NFH, neurofilament heavy polypeptide; PFC, prefrontal cortex; DPFC, dorsal prefrontal cortex; CC, corpus callosum; TL, temporal lobe; Hip, hippocampus; MT, mediodorsal thalamus; CSF, cerebrospinal fluid; SN, substantia nigra; LVN, lateral vestibular nucleus; MDD, major depressive disorder.

Besides, alpha-internexin was also found significantly downregulated average −3.9 times (−1.2 ~ −7.9) in the EVs of the PFC brain region. Alpha-internexin, is another neuronal intermediate filament protein, and three neurofilament subunits (NFL, NFM, MFH) have recently been identified as the pathological hallmarks of neuronal intermediate filament inclusion disease (NIFID) (89), a neurological disease of early onset with a variable clinical phenotype including frontotemporal dementia, pyramidal and extrapyramidal signs (90). The granin family protein secretogranin-2 (Scg2), which is the precursor for biologically active peptides such as neuropeptides, also exhibited significant differences (reduced average −4.5 times (−4.4 ~ −4.7)) in EVs of PFC between animal groups in our study. Indeed, other studies have shown that Scg2 is strongly modulated in cerebrospinal fluid (CSF) of AD patients, using a parallel reaction-monitoring mass spectrometric method (91). Scg2 was also identified and significantly changed in the CSF of multiple sclerosis patients (92). Additional neuropeptide precursor proteins ProSAAS (reduced average −5.5 times (−3.1 ~ −8.5)) and protachykinin-1 (Tac1, reduced average −3.7 times (−2.3 ~ −5.0)) also were identified significantly changed in the EVs of PFC. Previous immunoreactivity study has shown a novel function for ProSAAS protein as amyloid anti-aggregant in AD, which may play a role in AD pathology. Tac1 can be cleaved to 11 amino acids neuropeptide substance P, the first identified neuropeptide in the history, and corresponding receptor antagonist (aprepitant) has been developed approved by FDA for treatment of chemotherapy-induced emesis; it also has demonstrated clinical efficacy in the treatment of major depression (93). Furthermore, other neuropeptides precursors also identified in the EVs of brain regions such as Pro-MCH, vasopressin-neurophysin 2-copeptin (Avp), chromogranin-A, neuroendocrine protein 7B2.

## Discussion

This study employs a novel analytical approach to monitor EV associated proteins, in order to investigate the DBS regulation on different rat brain regions. This method has merits on high specificity, with the potential to uncover molecules linked to brain pathology. We discovered over a dozen of significantly changed proteins in PFC, hippocampus and striatum in DBS treatment animals compared with controls, some of which had been previously reported and linked to neurodegenerative diseases.

Neurofilaments (NF) are intermediate filament in neurons composed of three subunits, neurofilament light polypeptide (NFL), neurofilament medium polypeptide (NFM), and neurofilament heavy polypeptide (NFH). NF proteins are major components of large myelinated axons. Abnormal accumulation of NF proteins is a primary finding in several human neurodegenerative diseases, such as ALS, PD, AD and Charcot-Marie-tooth (CMT) (85). Brettschneider and colleagues reported that NFH protein increased 10 folds in CSF of AD patients compared with controls, using ELISA and antibodies (94). NFL is generally considered to be a biomarker of large-caliber myelinated axon injury (95) and in CSF as a biomarker of neurocytoskeleton impairment in AD (96). One paper has systematically reviewed that CSF NFL elevated average 2.35 (1.90 ~2.91) times in AD patients compared with controls (97). It suggested that the NEF was not only strongly associated with AD but also strongly associated with mild cognitive impairment due to AD. Besides, it also found that the NFL mRNA is selectively reduced by up to 70% in degenerating neurons of AD and ALS (98, 99). The proteomic analysis of substantia nigra (SN) showed reduced protein level of NFL and NFM in PD patients (100). NFL mRNA is also decreased and correlated with the severity of PD (101). In contrast, Abdo and colleagues (102) have reported significant increased CSF level of NFL and NFH in multiple system atrophy predominated by Parkinsonism. In our study, we found significant changed NFL and NFM in all of PFC, hippocampus and striatum in stimulated animals compared with controls, while NFH was not significantly modulated. The result shows that DBS brain stimulation can profoundly impact biomarker proteins that are sensitive indicators of neurodegenerative diseases.

Other CNS related diseases have also been linked to modulations of neurofilament proteins such as Schizophrenia (87), MDD (88) and suicide (86). We can conclude from the present study that EV associated NF proteins exhibit brain region specific regulation, and EV monitoring may thus provide a more sensitive opportunity to study the pathogenesis of related brain disorders.

We document that in all three brain regions, oxidative stress related proteins were found significantly changed in DBS stimulated animals compared to controls. High levels of oxidative stress have been implicated in many common neurodegenerative diseases. Neuron cells are particularly vulnerable to oxidative damage due to high polyunsaturated fatty acid content in membranes, high oxygen consumption, and weak antioxidant defense (84, 103). Neurodegenerative diseases are characterized by progressive damage in neural cells and neuronal loss, which lead to compromised motor or cognitive function. The overproduction of reactive oxygen species may have complex roles in promoting neurodegenerative diseases according to previous researches (104, 105). The exact molecular pathogenesis of this disturbance of redox balance remains unclear. Our findings certainly suggest that DBS brain stimulation might modulate or even reverse oxidative stress caused damage in CNS.

The protein SOD1 is thought to be an important cell-signaling molecule with neuromodulatory properties. Mutant SOD1 can contribute to the progression of familial ALS through the dysregulation of signal transduction pathways in motor neurons and in the activity of supportive glial cells (106, 107). Studies in vitro, CSF and in transgenic mouse models show that SOD1 is secreted via the microvesilcular secretory pathway(108–110). In our study, SOD1 was significant decreased average −2.3 times (−1.6 ~ −3.0) in stimulated PFC EVs related to control, demonstrating a strong association between DBS and CNS related diseases although the precise mechanism remains unclear. DNA damage process related proteins were also found frequently in our study through the functional analysis. Particularly, mitochondria dysfunction could be linked with sustained oxidative stress in neurodegenerative disorders, which would be have strongly relevant with DNA damage process.

## Conclusions

Brain stimulation profoundly impacts the regulation and transmission of signaling molecules in different brain regions. Here, we evaluate DBS effects on brain regions by MS-based EVs proteins quantitative analysis. Last, we got an amount of significant changed EVs proteins in PFC, hippocampus and striatum after stimulation on rat brain BF position. Through functional analysis, these proteins were involved in oxidative stress, nervous system development, DNA damage and protein biosynthetic processes, and some of them are strongly associated with CNS related disorders. Particularly, the NF proteins that were reported as potential biomarkers in many CNS related diseases. In our study, the NF subunits NFL and NFM were both significantly changed in EVs of PFC, hippocampus and striatum in stimulated samples compared with control. The strongly related to neurodegenerative disease SOD1 protein was also found significantly changed in PFC. All of this founding shown that the brain BF stimulation can impact the EVs proteins differential expression in different regions, which would be helping to explain the stimulation certainly potential effects on benefits of patients in the future.

This work is supported by the grants MOST 2015DFG424460, NSF 21475128 and CAS 100 Talent Project.

The authors declare no competing financial interest.

## Abbreviations

AD: Alzheimer disease;
ALS: amyotrophic lateral sclerosis;
BF: basal forebrain;
CID: collision induced disassociation;
CMT: Charcot-Marie-tooth;
CNS: central nervous system;
CSF: cerebrospinal fluid;
DBS: deep brain stimulation;
EV: extracellular vesicle;
FDA: federal drug & administrative
FDR: false discovery rate;
GABA: gamma-aminobutyric acid;
GPi: globus pallidus;
MCH: melanin-concentrating hormone;
MDD: major depressive disorder;
NBM: nucleus basalis magnocellularis;
NFH: neurofilament protein heavy peptide;
NFL: neurofilament protein light peptide;
NFM: neurofilament protein medium peptide;
NIFID: neuronal intermediate filament inclusion disease;
OCD: obsessive-compulsive disorder;
PD: Parkinson’s disease;
PFC: prefrontal cortex;
PTM: post-transnational modification;
SNpr: substantia nigra pars reticulata;
STN: subthalamic nucleus;
TEAB: triethylammonium bicarbonate;
TEM: transmission electromicropy;
VC: visual cortex;

